# Assessing the potential of environmental DNA metabarcoding for monitoring Neotropical mammals: a case study in the Amazon and Atlantic Forest, Brazil

**DOI:** 10.1101/750414

**Authors:** Naiara Guimarães Sales, Mariane da Cruz Kaizer, Ilaria Coscia, Joseph C. Perkins, Andrew Highlands, Jean P. Boubli, William E. Magnusson, Maria Nazareth Ferreira da Silva, Chiara Benvenuto, Allan D. McDevitt

## Abstract

The application of environmental DNA (eDNA) metabarcoding as a biomonitoring tool has greatly increased in the last decade. However, most studies have focused on aquatic macro-organisms in temperate areas (e.g., fishes). We apply eDNA metabarcoding to detect the mammalian community in two high-biodiversity regions of Brazil, the Amazon and Atlantic Forest. We identified critically endangered and endangered mammalian species in the Atlantic Forest and Amazon respectively and found congruence with species identified via camera trapping in the Atlantic Forest. In light of our results, we highlight the potential and challenges of eDNA monitoring for mammals in these high biodiverse areas.

## Introduction

A quarter of mammal species are endangered according to the IUCN Red List of Threatened Species (IUCN 2019) and there is clearly a need for more cost-effective and rapid methods for long-term biomonitoring to be applied across different biomes and over large spatial and temporal scales (Sales et al. 2019a). In recent years, environmental DNA (eDNA) metabarcoding (the simultaneous identification via next-generation sequencing of multiple taxa using DNA extracted from environmental samples, e.g., water, soil) has delivered on its initial potential and is now revolutionizing how we monitor biodiversity (Deiner et al. 2017). The majority of eDNA metabarcoding applications have focused on monitoring fish and macroinvertebrates, with mammals being targeted in only 8% of vertebrate studies (Tsuji et al. 2019). However, with the development of universal primers for vertebrates and mammals specifically, there has been a recent surge in studies tailored to detect and/or monitor mammalian communities in terrestrial and freshwater environments (e.g. Ushio et al. 2017, Harper et al. 2019, Sales et al. 2019a).

Several recent mammal-focused eDNA metabarcoding studies in temperate regions in the northern hemisphere have relied on well-studied systems with accompanying long-term or historical survey data to test the efficiency of this novel biomonitoring tool (e.g., Harper et al. 2019, Sales et al. 2019a). However, mammal conservation can be more challenging in biodiversity-rich countries as long-term monitoring systems are still scarce outside of Europe and North America (Proença et al. 2017) and ecological field studies, usually used to plug this gap, are often hindered due to difficulties of sampling over wide spatial scales. For effective conservation action, adequate knowledge regarding the biodiversity components present in each area is of paramount importance.

Environmental DNA from lentic and lotic systems has been found to be effective in not just monitoring aquatic and semi-aquatic mammals, but also terrestrial species (Harper et al. 2019, Sales et al. 2019a). Here, we explore the application of eDNA metabarcoding for Neotropical mammals by verifying its ability to detect aquatic and terrestrial animals from rivers/streams in the highly biodiverse biomes of the Brazilian Amazon and Atlantic Forest. The Amazon is the largest tropical rainforest on Earth, encompassing at least 10% of the world’s biodiversity. The Atlantic Forest, which is currently represented by only 11% of its original cover (Ribeiro et al. 2009), is the second most biodiverse biome in South America (WWF 2018).

## Methods

In the Amazon, water samples (500mL each, in three replicates) were obtained from six sites within three main areas (A-C; Fig. 1; Table S1). In the Atlantic Forest, water and sediment samples (500mL of water and 25mL of sediment, in three replicates) were obtained from eight sites located in two valleys of the Caparaó National Park (D-E; Fig. 1; Table S1). Temperature and pH were recorded at each site in the Amazon. Mammal-specific universal primers targeting the mitochondrial 12S rRNA gene were used (Ushio et al. 2017). The workflow was conducted following the protocol described in Sales et al. (2019a) and a more detailed description is included in the Supporting Information.

**Figure 1.**
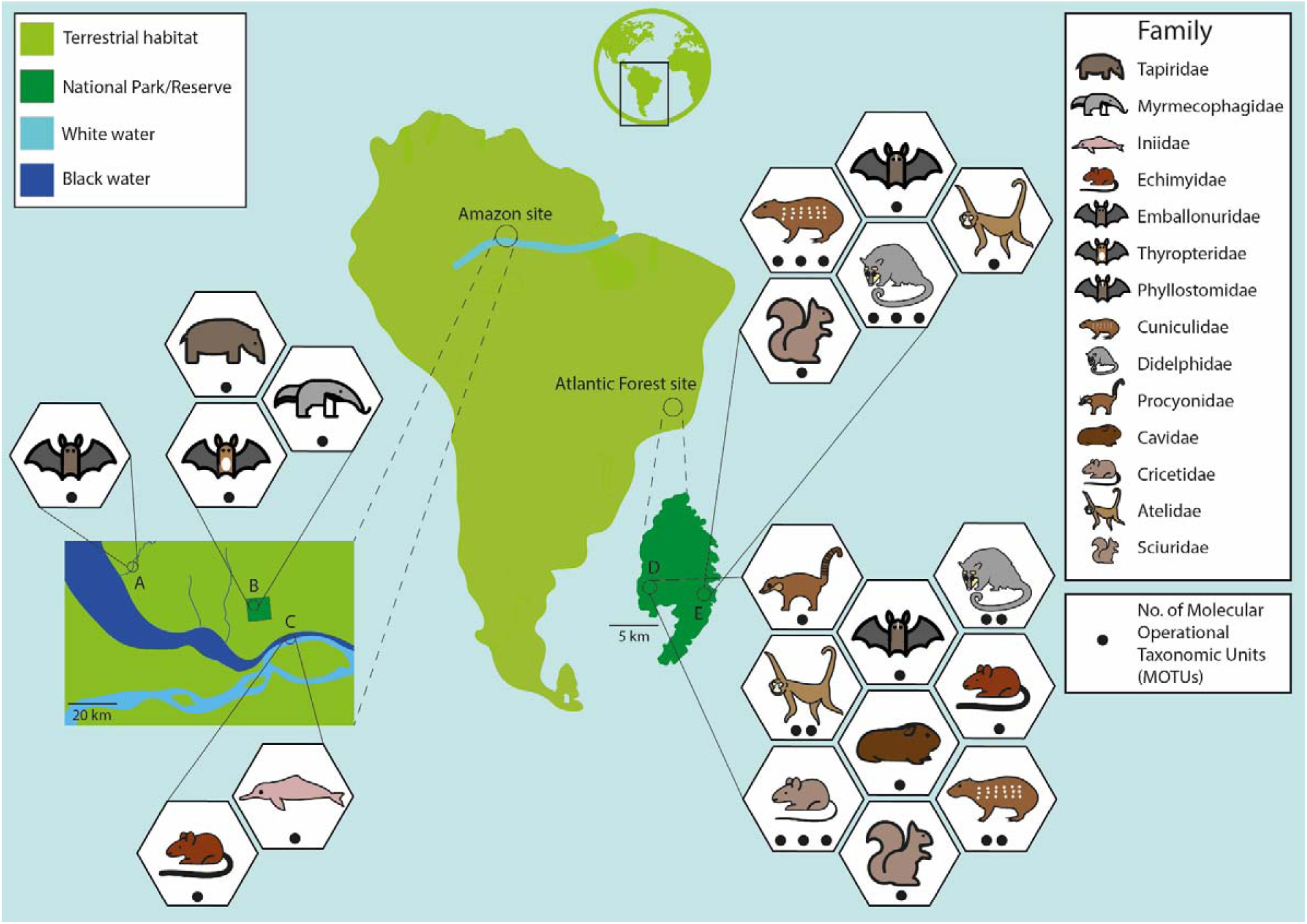
Sampling areas for environmental DNA (eDNA) in the Amazon (A-C) and Atlantic Forest (D-E) biomes in Brazil. The families recovered from eDNA metabarcoding in each area are represented by stylized drawings and the number of Molecular Operational Taxonomic Units (MOTUs) recovered within each family is indicated.

Additional data regarding species’ distribution in the Atlantic Forest were obtained through camera-trap surveys. Both valleys in the Caparaó National Park were surveyed with terrestrial and arboreal camera traps (Bushnell Trophy CamTM, USA; see Supporting Information).

## Results and Discussion

A total of ∼1.3 million mammal reads were obtained after all the bioinformatic filtering steps (Amazon – 833,623 reads; Caparaó – 109,233 reads for water samples and 334,593 for sediment samples). Only reads recovered for wild mammals (919,910 reads) were retained for downstream analyses.

Overall, we detected 28 Molecular Operational Taxonomic Units (MOTUs - Blaxter et al. 2005) from terrestrial and aquatic mammals, representing eight orders and 14 families (Table S2). Considering a threshold of >0.97 minimum identity, only 13 MOTUs could be assigned to the species level (Table S2). In the Amazon, six species were recovered, with three currently listed as endangered by the IUCN’s Red List (IUCN 2019) in different categories: the Endangered Amazon river dolphin (*Inia geoffrensis*), the Vulnerable giant anteater (*Mymercophaga tridactyla*) and the Vulnerable lowland tapir (*Tapirus terrestris*). Three Least Concern species were also identified: *Thyroptera discifera* and *Rhynchonycteris naso* in the order Chiroptera and the rodent *Toromys rhipidurus*. Only one MOTU was detected for each family (Fig. 1).

In Caparaó National Park, nine families were detected using eDNA: five in the west side of the park (D) and nine in the east side (E; Fig. 1 and S1). Of these, only seven could be assigned to the species level (Table S2). Here, camera-trap surveys detected 17 species (and additional unidentified small mammal species), encompassing 12 families (Fig. S2; Table S3). Combining the two non-invasive techniques, 15 families were detected overall (Table 1), six of which by both methods, three exclusively by eDNA metabarcoding and six solely by the camera traps. Although this study was not designed to provide a direct comparison between methods (e.g., Harper et al. 2019, Sales et al. 2019a), it highlights the potential of implementing multiple non-invasive approaches in providing an overview of the mammalian community composition in biodiversity rich areas.

**Table 1.** Number (*n*) of species captured with camera traps and number of Molecular Operational Taxonomic Units (MOTUs) captured with environmental DNA (eDNA) metabarcoding for orders and families within Caparaó National Park, Atlantic Forest. See Tables S2 and S3 for a more extensive breakdown of camera trap and eDNA data, respectively.

| Order | Family | Camera (*n* species) | eDNA (*n* MOTUs) |
| --- | --- | --- | --- |
| **Carnivora** | Felidae | 1 | - |
|  | Mustelidae | 1 | - |
|  | Procyonidae | 2 | 1 |
| **Chiroptera** | Phyllostomidae | - | 2 |
| **Didelphimorphia** | Didelphidae | 3 | 3 |
| **Pilosa** | Myrmecophagidae | 1 | - |
| **Primates** | Atelidae | 1 | 2 |
|  | Callithrichidae | 1 | - |
|  | Cebidae | 1 | - |
| **Rodentia** | Caviidae | 1 | 1 |
|  | Cricetidae | - | 3 |
|  | Cuniculidae | 1 | 3 |
|  | Echimyidae | 2 | 1 |
|  | Erethizontidae | 2 | - |
|  | Sciuridae | 1 | 1 |

More MOTUs were retrieved for the families detected in the Atlantic Forest, suggesting the occurrence of several species of the same family in this area. For example, three MOTUs were recovered in the east side and two from the west side of the Park for both Didelphidae and Cuniculidae. Camera trapping recorded three species of Didelphidae (*Caluromys philander, Didelphis* sp., *Philander frenatus*), in accordance with the eDNA data. Only one species from the Cuniculidae (*Cuniculus paca*) recorded by camera traps is known to occur in the Caparaó and the existence of three MOTUs for this family might be due to natural intraspecific genetic variability (Fig. 1). Cricetidae had three MOTUs in the west side of the Park: although this family was not identified by camera traps, several species are described for the Atlantic Forest, including endemic and recently described species (Gonçalves & Oliveira 2014). Furthermore, the Critically Endangered primate *Brachyteles hypoxanthus* was detected using eDNA, demonstrating its potential to detect arboreal mammals from water samples (Harper et al. 2019).

As a similar sampling effort was applied for both areas in this study, there is a need to consider what factors might explain the difference in the number of MOTUs recovered for each biome, particularly if we assume that mammalian alpha diversity should at least be as high in the Amazonian sampling sites as in the Caparaó forest site (see Costa et al. 2000). For example, all the families detected in the Atlantic forest that were not detected in the Amazonian samples are known to occur in Area B of the Amazon (Mendes Pontes et al. 2008). DNA degradation in water is one of the main factors reducing detectability over time and limiting temporal inferences. The sampled black waters in the Amazon have low pH (ranging from 3.85 to 4.27), whereas in the Caparaó the reported values are above 6.5 (Rodrigues 2015). Acidic environments show higher eDNA decay and lower persistence rate due to the increased degradation of DNA via chemical hydrolysis (Seymour et al. 2018). Therefore, the eDNA recovered in the low pH waters of the Amazon might be derived from specimens which have had very recent contact with the water body. Mammal eDNA recovery depends not only on the species presence but also on the direct/indirect contact with the water system (Harper et al. 2019). The junction of the Negro and Amazon Rivers (area C) has an enormous volume of water and possibly much time had elapsed since it flowed under the forest canopy, but the other Amazonian streams (area B; Fig. 1) are similar in size to those in the Atlantic Forest. In the Amazon, all species/MOTUs were detected in a single replicate, except for the lowland tapir (detected in four replicates in three different streams). This species is known to defecate more frequently in water than on land (Tobler et al. 2010) so this may explain its higher rates of eDNA detection. In the Atlantic Forest, several MOTUs/species were recovered from multiple replicates/sites (Fig. S1), suggesting longer persistence of eDNA in this environment.

There is a clear limitation in terms of available DNA sequences in public databases (e.g., Genbank) to either match identified MOTUs to species, or to distinguish between closely related species within the same genus. This issue has been highlighted in previous Neotropical eDNA studies for other taxonomic groups (Cilleros et al. 2019, Sales et al. 2019b). A 12S reference database exists for 164 Amazonian mammalian species in French Guiana (Kocher et al. 2017) and all Amazonian MOTUs were identified to species level here. However, this was not the case for the Atlantic Forest. This biome hosts more than 300 mammalian species (more than 50% of medium/large species considered at least Vulnerable; Souza et al. 2019). Therefore, for eDNA monitoring to be implemented in this biome, there is a clear need to generate reference DNA barcodes of a large proportion of the mammalian communities present.

Here, we demonstrated the potential of applying a cutting-edge non-invasive and cost-effective molecular approach for biodiversity assessment and systematic monitoring scheme of Neotropical mammals, including highly threatened species. This is particularly relevant given the current political climate in Brazil, which is resulting in research funding and environmental crises. However, significant challenges remain to implement this method in the Neotropics, from a better understanding of the ecology of eDNA within these variable environments, to the current lack of appropriate reference sequences for species determination in these biodiversity-rich and anthropogenically-impacted biomes.

## Acknowledgments

This project was partially funded by the University of Salford Internal Research Award awarded to CB, ADM and IC. We are grateful to Vitor Borges for assistance in the field. The present study was carried out with all required permits (ICMBIO N. 54795-2, DEFRA 126191/385550/0).

## Author Contributions

NGS, MDCK, ADM, CB, IC, JPB, WEM and MNFS conceived, and NGS, MDCK, ADM, IC and CB designed the study. NGS, MDCK, AH and CB carried out the eDNA sampling. NGS, MDCK and JCP performed the laboratory work. NGS carried out the bioinformatic analyses. MDCK analysed the camera trap data. NGS, ADM and IC analysed the eDNA data. NGS, ADM and MDCK wrote the paper, with all authors contributing to editing and discussions.

