## Supplemental Methods, Tables S1 and S3 and Figs. S1 and S2 for "Assessing the potential of environmental DNA metabarcoding for monitoring Neotropical mammals: a case study in the Amazon and Atlantic Forest, Brazil"

**Material and Methods**

*eDNA sampling*

The present study was carried out with all required permits (ICMBIO N. 54795-2, DEFRA 126191/385550/0). In the Brazilian Amazon, a total of 41 samples were obtained from three main areas in close proximity in Amazonas State (Fig. 1) in January 2019. Sampling location A was in the Rio Aturiá, a tributary leading into the Rio Negro (two replicates). Sampling location B consisted of four streams within the Adolpho Ducke Reserve, Manaus (with nine replicates taken in each stream). Sampling areas A and B are blackwater. Sampling location C was the ‘Meeting of the Waters’, the blackwater from the Rio Negro and the whitewater of the Solimões form the Amazon River (three replicates). A total of four field blanks were taken.

In the Atlantic Forest, a total of 48 samples (24 water samples and 24 sediment samples) were collected in February 2018 from eight sites located in two different areas (Aleixo Valley (D) and Santa Marta Valley (E); Fig 1) in the Caparaó National Park, State of Minas Gerais, Brazil. In each sample site, three replicates of each sampling medium were collected from three streams and one main river. A total of two field blanks were taken.

The same water collection protocol was used for both sampling areas, and sediment samples were obtained only from the Caparaó National Park. Water samples were collected using sterile bottles of 500 mL each and filtered using 50mL sterile syringes (Thermo Co., Tokyo, Japan) and 0.45μm Sterivex capsule filters (Millipore, MA, USA) immediately after sampling. Filters were stored in silica beads at room temperature until DNA extraction. Sediment samples (~25mL) were collected from the superficial layer using centrifuge tubes (50mL) following the same sampling approach adopted for water samples (three replicates obtained from the same sample sites where water samples were collected). Sediment samples were stored in 100% ethanol and kept at room temperature until DNA extraction.

*Laboratory work*

eDNA was extracted following an adapted Mu-DNA protocol for both water and sediment samples (Sellers et al. 2018). An extraction blank was included for each batch of samples processed. Amplification of the mitochondrial 12S rRNA gene fragment (˜171 bp) was carried out in triplicate on all eDNA samples and negative control samples using the MiMammal set of primers (previously described for metabarcoding mammalian eDNA – Ushio et al., 2017). Library preparation was conducted according to the same protocol described in Sales et al. (2019). Adequate decontamination precautions were taken to reduce the risk of contamination between samples and to verify the occurrence of potential contamination field and laboratory blanks (sample collection, DNA extraction, PCR) were obtained and processed alongside collected samples. A total of 108 samples (including field, DNA extraction and PCR blanks) were sequenced in two multiplexed Illumina MiSeq (www.illumina.com) runs alongside with samples belonging to non-related projects.

*Bioinformatics analyses*

The OBITools metabarcoding package (Boyer et al., 2016) was used for the bioinformatics analysis following the approach described in Sales et al. (2019). In order to remove false positives and MOTUs likely originating from contaminations or sequencing errors we applied the following framework: i) the maximum number of reads detected in the controls was removed for each MOTU from all samples; ii) MOTUs containing less than 10 reads were discarded; iii) obvious non-target species or those MOTUs likely originating from carry-over contaminations were removed from the dataset (Li et al., 2018; Ushio et al., 2018). For the taxonomic assignment, after these stringent filtering steps, a total of 28 MOTUs (excluding humans and domestic mammals) were retained and a threshold of sequence identity of at least 97% was used to assign the MOTU at species level (N=12). MOTUs showing similarities below the 0.97 minimum identity cut-off were considered only up to family level (see Table S2).

***Camera trapping***

Both sites in the Caparaó National Park were surveyed with terrestrial and arboreal camera trapping (Bushnell Trophy Cam^TM^, Bushnell Outdoor Products, USA) as part of ongoing monitoring projects (M. Kaizer et al., *unpublished data*).

*Ground camera trapping* - camera traps were deployed randomly in each site as part of an ongoing study on terrestrial mammals’ inventory (M. Kaizer et al., *unpublished data*). In the D Valley (see Fig. 1), one camera was deployed at three distinct and non-overlapping sampling points in 2017 (March and August) and 2018 (March to November) for a total of 121 camera trapping days. Likewise, one camera was deployed at two distinct sampling points in the E Valley in February 2016 and May to October 2018, totaling 168 camera trapping days. Cameras were attached to a tree at approximately 50 cm above ground using a cable lock to avoid thefts, and no baits or lures were used. Cameras were set to work 24h/day, at normal sensitivity, and at two shot bursts followed by a 30 second video.

*Arboreal camera trapping* - eight camera traps were deployed simultaneously in the forest canopy from January 2017 in the E Valley and February 2017 in the D Valley as part of an ongoing study on primate monitoring (M. Kaizer et al., *unpublished data*). Arboreal cameras were attached to a tree at a mean height of 11 m above ground and an average distance of 255.7 m ± 66.9 (SD) apart in each site. Cameras were set to be active continuously, at a hybrid mode (two pictures followed by a 30 second video), and a 10 second interval between triggers. These cameras were in operation until December 2017, resulting in an effort of 1538 and 1305 camera trapping days at the D and E Valleys, respectively.

Table S1. Coordinates and dates of eDNA sampling localities in the Atlantic Forest and Amazon respectively. Information is provided on which samples were placed on each of two MiSeq sequencing runs.

| **eDNA Samples** | | | | | |
| --- | --- | --- | --- | --- | --- |
| **Caparaó National Park** | | | | | |
| **Samples** | **Locality** | **Fig. 1 Map** | **Lat** | **Long** | **Date_Sampling** |
| C1 | Vale do Aleixo_Field Blank | - | - | - | 26.02.18 |
| C2 | Santa Marta_Field Blank | - | - | - | 27.02.18 |
| EB1 | Extraction Blank_water | - | - | - | - |
| EB2 | Extraction Blank_water | - | - | - | - |
| EB2 | Extraction Blank_sediment | - | - | - | - |
| 1_1-1_3 | Vale do Aleixo | D | -20.48004082 | -41.84522942 | 26.02.18 |
| 2_1-2_3 | Vale do Aleixo | D | -20.48052212 | -41.84725972 | 26.02.18 |
| 3_1-3_3 | Vale do Aleixo | D | -20.48208208 | -41.8517721 | 26.02.18 |
| 4_1-4_3 | Vale do Aleixo | D | -20.48247262 | -41.85437558 | 26.02.18 |
| 5_1-5_3 | Santa Marta | E | -20.49156406 | -41.75546539 | 27.02.18 |
| 6_1-6_3 | Santa Marta | E | -20.49142451 | -41.75515629 | 27.02.18 |
| 7_1-7_3 | Santa Marta | E | -20.49034586 | -41.75087329 | 27.02.18 |
| 8_1-8_3 | Santa Marta | E | -20.49017437 | -41.74345422 | 27.02.18 |
|  |  |  |  | Water samples | 24 |
|  |  |  |  | Sediment Samples | 24 |
|  |  |  |  | Collection Blanks | 2 |
|  |  |  |  | Extraction Blanks | 3 |
| **Amazon** | | | | | |
| **Sample** | **Locality** | **Fig. 1 Map** | **Lat** | **Long** | **Date_Sampling** |
| **DK01** | BLANK_Acará_L2 |  | - | - | 18.01.19 |
| **DK02** | BLANK_Acará_L4 |  | - | - | 17.01.19 |
| **DK03-DK11** | Acará_L4 | B | -2.95083333 | -59.95694444 | 17.01.19 |
| **DK12-DK20** | Acará_L2 | B | -2.93611111 | -59.96277778 | 18.01.19 |
| **DK21** | Barro Branco | B | -2.92638889 | -59.97027778 | 15.01.19 |
| **DK22** | BLANK_extraction |  | - | - | - |
| **DK23** | BLANK_Acará_L3 |  | - | - | 16.01.19 |
| **DK24** | BLANK_Barro Branco |  | - | - | 15.01.19 |
| **DK25-DK26** | Barro Branco | B | -2.92638889 | -59.97027778 | 15.01.19 |
| **DK27-DK28** | Aturiá | A | -2.03583333 | -61.14944444 | 21.01.19 |
| **DK29-DK34** | Barro Branco | B | -2.92638889 | -59.97027778 | 15.01.19 |
| **DK35-DK43** | Acará_L3 | B | -2.94472222 | -59.955 | 16.01.19 |
| **DK44-DK46** | Solimões | C | -3.12888889 | -59.89638889 | 26.01.19 |
| **DK47** | BLANK_extraction |  | - | - | - |
|  |  |  |  | Water samples | 41 |
|  |  |  |  | Collection Blanks | 4 |
|  |  |  |  | Extraction Blanks | 2 |
| **RUN 1** |  |  |  |  |  |
| Caparao Water (24 samples + 2 collection blanks + 2 DNA extraction blanks + 4 PCR blanks) | | | | | |
| **RUN 2** |  |  |  |  |  |
| Amazon (41 samples + 4 collection blanks + 2 DNA extraction blanks + 4 PCR blanks) | | | | | |
| Caparao sediments (24 samples + 1 DNA extraction blank) | | | | | |

Table S2. Identified Molecular Operational Taxonomic Units (MOTUs) and their assignment to family, genus and species levels (Table as a separate .xlsx file).

Table S3. Species captured on ground (G) and canopy (C) camera traps in Caparaó National Park, Atlantic Forest.

| **Order** | **Family** | **Genus** | **Species** | **Valley D** | **Valley E** |
| --- | --- | --- | --- | --- | --- |
| **Carnivore** | Felidae | *Leopardus* | *Leopardus guttulus* | G | C \| G |
|  | Mustelidae | *Eira* | *Eira barbara* | G | C |
|  | Procyonidae | *Nasua* | *Nasua nasua* | G | C \| G |
|  |  | *Potos* | *Potos flavus* | - | C |
| **Chiroptera** | - | - | - | - | C |
| **Didelphimorphia** | Didelphidae | *Caluromys* | *Caluromys philander* | C | C |
|  |  | *Didelphis* | *Didelphis sp.* | - | C |
|  |  | *Philander* | *Philander frenatus* | - | C |
| **Pilosa** | Myrmecophagidae | *Tamandua* | *Tamandua tetradactyla* | - | C |
| **Primates** | Atelidae | *Brachyteles* | *Brachyteles hypoxanthus* | C | C |
|  | Callithrichidae | *Callithrix* | *Callithrix flaviceps* | C | C |
|  | Cebidae | *Sapajus* | *Sapajus nigritus* | C | C |
| **Rodentia** | Cuniculidae | *Cuniculus* | *Cuniculus paca* | - | G |
|  | Echimyidae | *Phyllomys* | *Phyllomys sp.* | C | - |
|  |  | *Trinomys* | *Trinomys sp.* | - | G |
|  | Erethizontidae | *Coendou* | *Coendou spinosus* | C | C |
|  |  | *Coendou* | *Coendou prehensilis* | C | - |
|  | Sciuridae | *Sciurus* | *Sciurus ingrami* | - | C \| G |
| **Unidentified small mammals** | - | - | - | C | C \| G |

Figure S1. Bubble graph representing presence-absence and categorical values of the number of reads retained (after bioinformatic filtering) for eDNA (water in blue and sediment in orange) from each family identified in each site (Valleys D and E) in Caparaó National Park, Atlantic Forest.

Figure S2. Collage of images representing examples of mammals captured from ground and canopy camera traps in Caparaó National Park, Atlantic Forest. Red boxes are used to highlight where the smaller mammals are positioned within the photograph. A: *Brachyteles hypoxanthus*; B: *Coendou spinos*; C: *Cuniculus paca*; D: *Nasua nasua*; E: *Sciurus ingrami* and F: *Phyllomys* sp.

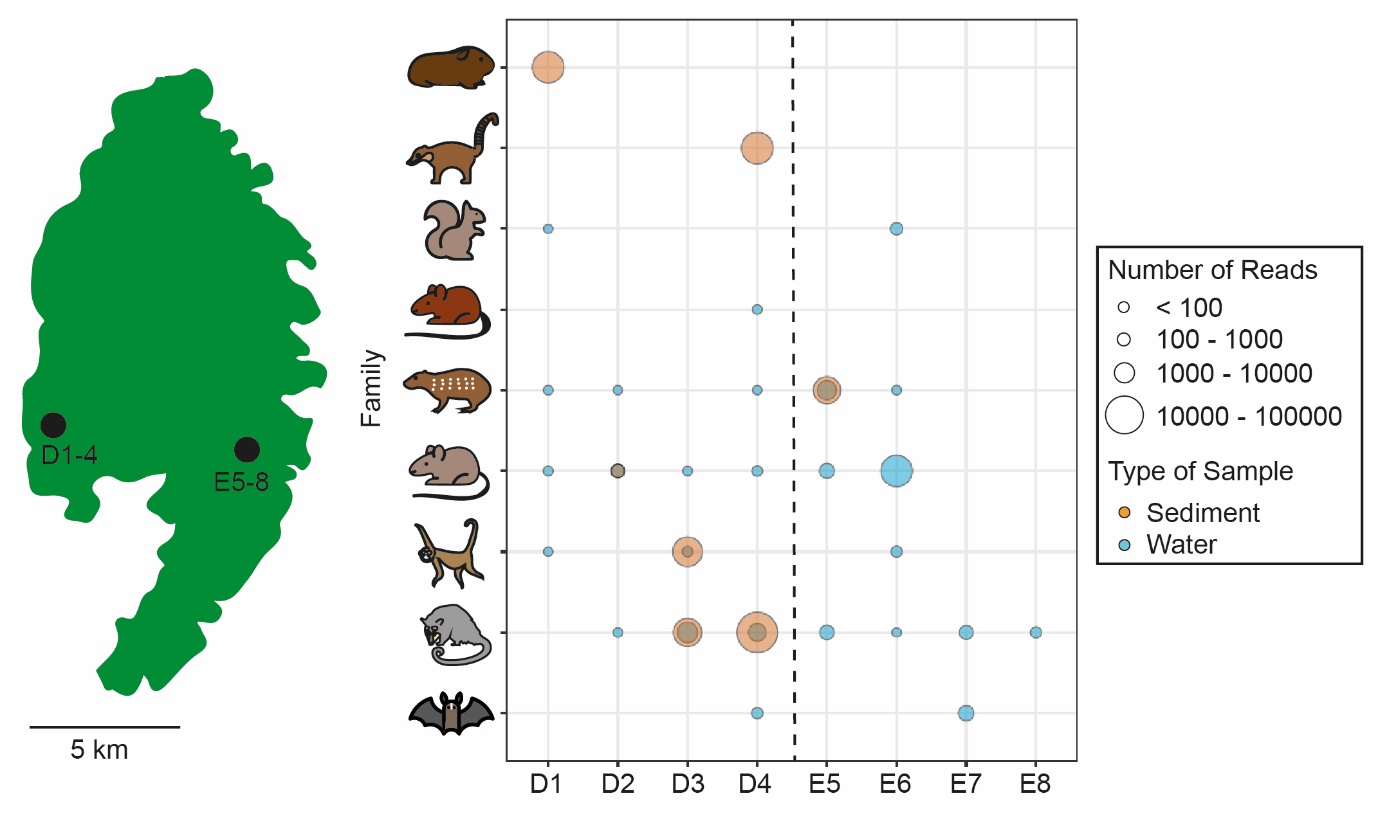

Figure S1.

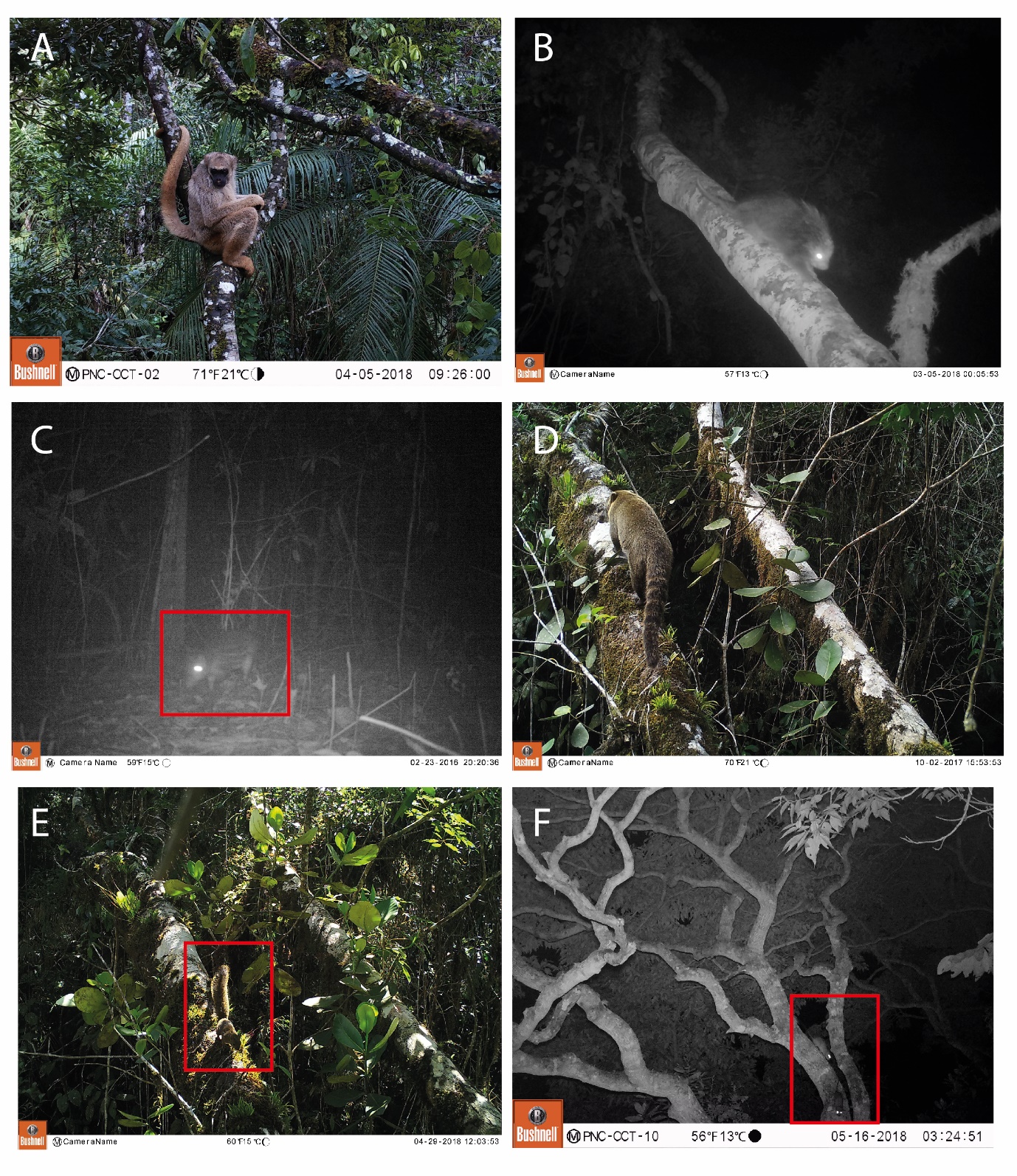

Figure S2.
